# Mechanism Driven Early Stage Identification and Avoidance of Antisense Oligonucleotides Causing TRL9 Mediated Inflammatory Responses in Bjab cells

**DOI:** 10.1101/2021.12.12.472280

**Authors:** Adam J. Pollak, Patrick Cauntay, Todd Machemer, Suzanne Paz, Sagar Damle, Scott P. Henry, Sebastien A. Burel

## Abstract

Nucleic acid-based PS-ASOs have the potential to activate cellular innate immune responses, and the level of activation can vary quite dramatically with sequence. Minimizing degree of proinflammatory effect is one of the main selection criteria for compounds intended to move into clinical trials. While a recently developed hPBMC-based assay showed excellent ability to detect innate immune active PS-ASOs, which can then be discarded from the developmental process, this assay is highly resource-intensive and easily affected by subject variability. This compelled us to develop of a more convenient high-throughput assay. Here, we describe a new in vitro assay, utilizing a cultured human Bjab cell line, which was developed and validated to identify PS-ASOs that may cause innate immune activation. The assay was calibrated to replicate results from the hPMBC assay. The Bjab assay was designed to be high throughput and more convenient by using RT-qPCR readout of mRNA of the chemokine Ccl22. The Bjab assay was also shown to be highly reproducible and to provide a large dynamic range in determining the immune potential of PS-ASOs via comparison to known benchmark PS-ASO controls that were previously shown to be either safe or inflammatory in clinical trials. In addition, we demonstrate that Bjab cells can be used to provide mechanistic information on PS-ASO-TLR9 dependent innate immune activation.

## INTRODUCTION

Phosphorothioate containing antisense oligonucleotides (PS-ASOs) are short, single stranded nucleic acids that bind to specified RNAs via Watson-Crick base pairing to modulate corresponding protein levels (Crooke, Baker et al. 2021). Various changes to PS-ASO backbone and 2’ modifications have been employed to enhance PS-ASO stability and potency while also mitigating potential toxicities (Crooke, Liang et al. 2021). PS-ASO design is evolving to maximize their therapeutic index and to increase the ways in which they can modulate cellular RNAs (Anderson, Freestone et al. 2021, Crooke, Liang et al. 2021). Several PS-ASOs are either approved to be used as drugs or are in late-stages of clinical trials (Benson, Waddington-Cruz et al. 2018, Bennett, Krainer et al. 2019). In addition to the intended pharmacology, PS-ASOs have been found to cause unwanted effects, including innate immune activation of B cells, dendritic cells, or macrophages (Monteith, Henry et al. 1997, Burel, Machemer et al. 2012, Frazier 2015).

Previous reports suggest that PS-ASOs cause innate immune responses through Toll like receptor 9 (TLR9), which is a cell-surface and endosomal nucleic acid receptor (Senn, Burel et al. 2005, Paz, Hsiao et al. 2017). TLR9 is canonically considered a receptor to single or double stranded oligonucleotides (ODNs) that must contain at least one unmethylated CpG site (Lind, Rael et al. 2021). These CpG sites are underrepresented in mammalian DNA yet are present in microbial species (Lind, Rael et al. 2021). Thus, TLR9 is thought to be only activated when the cell encounters evidence of microbial DNA.

Interestingly, it has been shown that PS-ASOs without unmethylated CpG sites can activate innate immune responses, albeit with much less potency for activation than sequences including canonical unmethylated CpG sites (Younis, Vickers et al. 2006, Paz, Hsiao et al. 2017). Such activation is mostly attributed to TLR9 (Vollmer, Weeratna et al. 2004, Paz, Hsiao et al. 2017) although there is evidence of the involvement of other innate immune receptors for some PS-ASOs (Burel, Machemer et al. 2012). TLR9-dependent responses are highly sequence dependent and only a few select sequences have been reported to produce immune responses (Paz, Hsiao et al. 2017). Mouse models have been used extensively to rank order PS-ASO immune responses with the goal of identifying features that minimize the inflammatory effects of PS-ASOs (Paz, Hsiao et al. 2017). However, there are considerable differences between species in the expression profile of TLR9 and TLR9’s propensity for interacting with certain ligands, and this has been well documented when comparing humans and mice (Pohar, Yamamoto et al. 2017). Therefore, it is not surprising that sequences that are not inflammatory in mice but are inflammatory in humans have been reported (PMBC). Even in rodents, there can be considerable differences in the magnitude of inflammatory response induced by non-CpG ODNs in a sequence- and TLR9-dependent (Paz, Hsiao et al. 2017). As such, in vivo mouse studies are used extensive to rank-order the level of innate immune stimulation with a goal of minimizing the inflammatory effects of ASOs. Nonetheless, in one example, ISIS 353512 (which does not have an unmethylated CpG site) caused evidence of innate immune activation in healthy trial volunteers. This PS-ASO did not indicate any level of potential for causing immune activation in non-human preclinical models (Szalai, McCrory et al. 2014). Exactly how PS-ASO without CpG sites activate human TLR9 remains incompletely understood yet it is clear that such effects are not necessarily replicated in animal models. Therefore, we hypothesized that despite the absence of canonical CpG motif, some ASOs such as ISIS 353512 might be inducing a TLR9 mediated response involving primarily B cells (Vollmer, Janosch et al. 2002, Elias, Flo et al. 2003, Vollmer, Weeratna et al. 2004) and that the lack of ISIS 353512 proinflammatory stimulation in rodent may stem from species-specific recognition of CpG and non-CpG ODN (Campbell, Cho et al. 2009). In an effort to avoid PS-ASOs with the potential to activate human TLR9 from entering clinical development, a human peripheral blood mononuclear cell (PBMC)-based in vitro assay was recently developed (Burel, Machemer et al. 2021). Using information from clinical trial data to provide benchmark PS-ASOs as controls, hPBMCs were shown to reliable recapitulate human innate immune responses to PS-ASOs using Il-6 protein production as a readout. Unfortunately, this assay has some drawbacks. Namely, the hPBMC assay requires the cumbersome process of collecting fresh PBMC from donors. In addition to being resource intensive, this introduces donor-to-donor genetic variability to the assay that is reflected in the varied response to the positive control, ISIS 353512. Therefore, we sought to develop a convenient, reproduceable, and high throughput cell-line based assay to test PS-ASO innate immune responses.

This paper describes our efforts to establish an in vitro model using an established cell line to improve throughput and reproducibility compared to human PBMCs. Following identification of a suitable cell-line among several candidates, head-to-head comparisons were made between the in vitro assay platforms to cross -validate the new model and to establish its read-out parameters. Furthermore, the endpoints were optimized to accommodate more efficient analysis for the new model. This new cell line-based model also proved useful by providing a convenient platform for better understanding the mechanism of innate immune cell activation by non-CpG ODNs.

## MATERIAL AND METHOD

### ODN design, synthesis and preparation

All oligonucleotides were designed and synthesized at Ionis Pharmaceuticals (supplemental table 1). To identify mouse ASO inhibitors, rapid throughput screens were performed in vitro as previously described (Watts, Manchem et al. 2005). The first 3 to 5 bases and last 3 to 5 bases of chimeric ASO have a 2’-O-(2-methoxy)-ethyl modification, and the ASOs also have a phosphorothioate backbone. ASOs were diluted in phosphate-buffered saline (PBS) for both in vivo and in vitro usage.

### Human peripheral blood mononuclear cells (PBMC) assay

Whole blood was collected from human volunteer donors with informed consent in 8 to 10 BD Vacutainer CPT 8 ml tubes (BD Biosciences, San Jose, CA). The blood samples were mixed immediately prior to centrifugation by gently inverting tubes 8-10 times. The CPT tubes were centrifuged at room temperature in a horizontal (swing-out) rotor for 30 min. at 1500-1800 RCF with brake off. The PBMC were retrieved at the interface between Ficoll and polymer gel layers and transfer to a sterile 50 ml conical tube; pooling up to 5 CPT tubes / 50 ml conical tube / donor. The cells were washed twice with PBS (Ca^++^, Mg^++^free; GIBCO) prior to another round of centrifugation. The supernatant was discarded without disturbing pellet. The cells were resuspended in RPMI1640+10% FBS+ penicillin /streptomycin. The cell density was estimated, and the density adjusted as to plate cells at 5 x 10^5^ in 50 μl/well of 96-well, sterile, round bottom, polypropylene plates for each ODN concentration tested for each donor. The ODNs 1:5 serial dilutions [2×] prepared in medium (RPMI1640+10% FBS+ pen/strep) in a separate sterile, V-bottom, polypropylene plate starting with the highest concentration (400 μM) in the top row.

Added 50 ml/well of 2x concentrated ODN diluted in RPMI1640+10% FBS+penicillin /streptomycin. The plated cells were incubated for 24 hours at 37°C; 5% CO_2_. Plates centrifuged at 330 x g for 5 minutes before removing supernatants to a separate sterile polypropylene 96-well plate. Cell supernatants stored at −70°C until processing for cytokine assay profiling.

### Cell Lines

The following cell lines were obtained from DSMZ cell repository (Braunschweig, Germany): Bjab (ACC 757), MEC2 (ACC 500), Daudi (ACC 78), KMS12BM (ACC 551). The following cell lines were obtained from ATCC (Manassas, VA): RPMI8226 (CCL-155), Db (CRL-2289), Jeko1 (CRL-3006), Rec1 (CRL-3004), EB1 (HTB-60), EB2 (HTB-61), EB3 (CCL-85), SUDHL16 (CRL-2964), Daudi (CCL-213), SUDHL5 (CRL-2958). KARPAS422 cells (06101702) were obtained from Millipore Sigma (St Louis, MO). The various cell lines were culture in RPMI1640+20% FBS+ penicillin /streptomycin. The cell density was set at 5 x 10^5^ in 50 μl/well of 96-well, sterile, round bottom, polypropylene plates for each ODN concentration tested for each donor. The ODNs 1:5 serial dilutions [2x] prepared in medium (RPMI1640+20% FBS+ pen/strep) in a separate sterile, V-bottom, polypropylene plate starting with the highest concentration (100 μM) in the top row. Added 50 μl / well of 2x concentrated ODN diluted in RPMI1640+20% FBS+penicillin/streptomycin. The plated cells were incubated for 24 hours at 37 C; 5% CO2. Plates centrifuged at 330 x g for 5 minutes before removing supernatants to a separate sterile polypropylene 96-well plate. Cell supernatants stored at −70°C until processing for cytokine assay profiling. RNA was isolated as described in later sections.

### Meso Scale Discovery platform

Multiplex plates pre-coated with capture antibodies for specific cytokines, along with diluents were brought to room temperature prior to use. Diluted standards (3 replicates) and undiluted samples were added to the wells and incubated for two hours in the Meso Scale Discovery^®^ Custom array. All incubations were performed at room temperature with vigorous shaking (300-1000 rpm). After the two-hour incubation, the plate was decanted and washed three times before adding a cocktail of Sulfo-Tag^®^ detection antibodies to each well. After incubating with detection antibodies for two hours, the plate was again decanted and washed three times. Read buffer was added to the wells using reverse pipetting techniques to ensure bubbles are not created to interfere with imaging. The plate was immediately imaged using the MSD Sector 2400 imaging system, and data was analyzed using MSD Discovery Workbench^®^ software. Concentrations of all unknown samples were back-calculated using results interpolated from the corresponding standard curve regression using a weighted, four-parameter fit. Final sample concentrations (pg/ml) were calculated by factoring the dilution factor used for each sample.

### RNA preparation and qRT-PCR

Total RNA was prepared using a RNeasy mini kit (Qiagen) from cells grown in 96-well plates using the manufacturer’s protocol. qRT-PCR was performed in triplicate using TaqMan primer probe sets. Briefly, ~50 ng total RNA in 5 μl water was mixed with 0.3 μl primer probe sets containing forward and reverse primers (10 μM of each) and fluorescently labeled probe (3 μM), 0.5 μl RT enzyme mix (Qiagen), 4.2 μl RNase-free water, and 10 μl of 2 × polymerase chain (PCR) buffer in a 20 μl reaction. Reverse transcription was performed at 48°C for 10 min, followed by 94°C for 10 min, and then 40 cycles of PCR were conducted at 94°C for 30 s, and 60°C for 30 s within each cycle using the StepOne Plus RT-PCR system (Applied Biosystems). The mRNA levels were normalized to the amount of total RNA present in each reaction as determined for duplicate RNA samples using the Ribogreen assay (Life Technologies). Ccl22 mRNA level were measured using the following primer probe set: Forward primer probe: CGCGTGGTGAAACACTTCTA, reverse primer probe: GATCGGCACAGATCTCCTTATC, detection probe: TGGCGTGGTGTTGCTAACCTTCAX. GAPDH mRNA level were measured and used as housekeeping gene for internal normalization using the following primer probe set: Forward primer probe: GAAGGTGAAGGTCGGAGTC, reverse primer probe: GAAGATGGTGATGGGATTTC, detection probe: CAAGCTTCCCGTTCTCAGCC.

### RNA sequencing

RNA was extracted from pulverized Bjab cells using Lysing matrix D beads (MP Biomedicals) and standard TRIzol extraction (ThermoFisher scientific). RNA quality was determined using RNA Pico Chips on Bioanalyzer 2100 (Agilent) and 3’ mRNA libraries (Lexogen, QuantSeq 3’ Digital Gene Expression (DGE)) were generated from high quality total RNA (RNA integrity number >8.0). Samples were subjected to single end sequencing on a NovaSeq platform (Illumina). Reads were aligned to mouse reference genome (mm10) using STAR 2.5.1b and ensembl build 96 gene models. Read count analysis was performed using salmon 0.7 (Patro, Duggal et al. 2017). Transcripts of interest were identified using JMP 16 (SAS Institute Inc, Cary, NC) by performing a linear correlation between the Log10 of TNF-a as measured by MSD with the Log2 TPM for each transcript detected by DGE. Rsquare for the top 47 transcripts is shown in supplemental table 1. Pathways were analyzed using Enrichr and the Kyoto Encyclopedia of Genes and Genomes database (Chen, Tan et al. 2013, Kuleshov, Jones et al. 2016, Xie, Bailey et al. 2021).

## RESULTS

### Selection of a cell line that can discriminate between low and highly inflammatory PS-ASO sequences

We sought to identify a cell line which could recapitulate the range of responses to PS-ASOs achieved using the validated hPBMC assay, which would offer a more robust and reproducible means of screening PS-ASOs. Previous evidence suggests that TLR9 was the most likely receptor mediating the inflammatory response to PS-ASOs (Paz, Hsiao et al. 2017) (Paz et al, and others). Therefore, we sought to query several cell lines which exhibited high levels of TLR9 mRNA. Expression data for TLR9 mRNA were obtained from 2 databases (CCLE (Barretina, Caponigro et al. 2012) or GENT (Shin, Kang et al. 2011)). Some cell lines such as Bjab were only present in one of the databases (listed as NA in the figure 1). TLR9 mRNA expression was not confirmed independently and TLR9 protein was not determined. The presence of TLR9 mRNA expression in the cell lines was highest among B cell derived lymphoma cell lines such as Burkitt Lymphoma and diffuse large B cell lymphomas likely compared to the normal expression of TLR9 in human B cells (Figure 1) (Avalos and Ploegh 2011).

**Figure 1:**
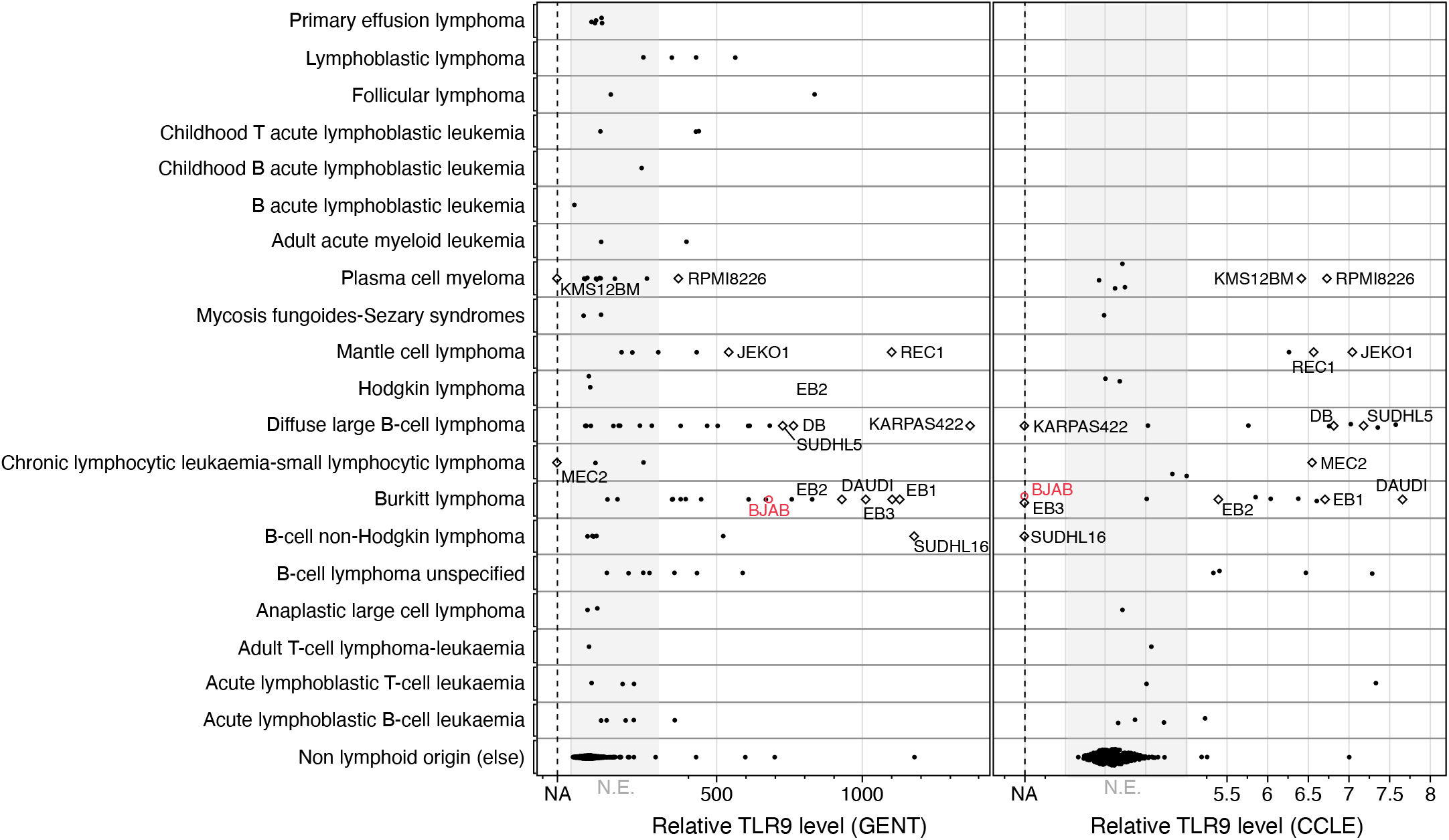
Relative TLR9 expression of a compendium of cell lines reported in 2 public databases (CCLE and GENT). Among cell lines characterized, TLR9 Expression is mostly limited to a subset of lymphomas. On X axis, NA represent the absence of reported data in either database. Cell lines selected for screening are indicated by a diamond. Cell lines of various origins are reported as ‘non_lymphoid_origin’. Bjab cells are highlighted in red. N.E. = Not expressed

We selected 14 cell lines with high reported expression in TLR9 mRNA in either CCLE or GENT (Figure 1) and assessed cytokine or chemokine production in response to stimulation with either control ISIS 104838 (minimally inflammatory Burel et al PBMC paper), or the more inflammatory ISIS 353512, ISIS 518477 or ISIS 120704 sequences. ISIS 120704 contains nonmethylated Type B CpG motifs and serves as a positive control for TLR9 activation (Pohar, Kuznik Krajnik et al. 2015). Our primary objective was to identify a cell line for which at least one analyte could discriminate between the benchmark ISIS 104838 and ISIS 353512 to a similar degree that was achieved with hPBMC cells. We measured levels of inflammatory cytokines and chemokines by MSD multiplexing (Figure 2, Supplemental Figure S1). The overall response of those various cell lines to the 3 more proinflammatory ODNs was very heterogenous. This is well illustrated by the broad range of response to ISIS 120704, the most proinflammatory ODN, for which some cell lines such a KARPAS422, MEC2, and SUDHL5 showed very little increase in TNF-α, IL-6, IL-10 or MIP-1B production (Supplemental Figure S1). In cell lines in which ISIS 120704 induced a response, the nature and magnitude of the analytes increased was variable with some cell lines such as EB1, EB2 or REC1 exhibiting strong IL-6 production, yet all three showed minimal increase in IL-10 production while only Rec1 increased TNF-α production in conjunction with IL-6. The responses to the inflammatory PS-ASOs without CpG sites was even more erratic, by both comparing responses between cell lines and by comparing responses of different analytes from the same cell lines.

**Figure 2.**
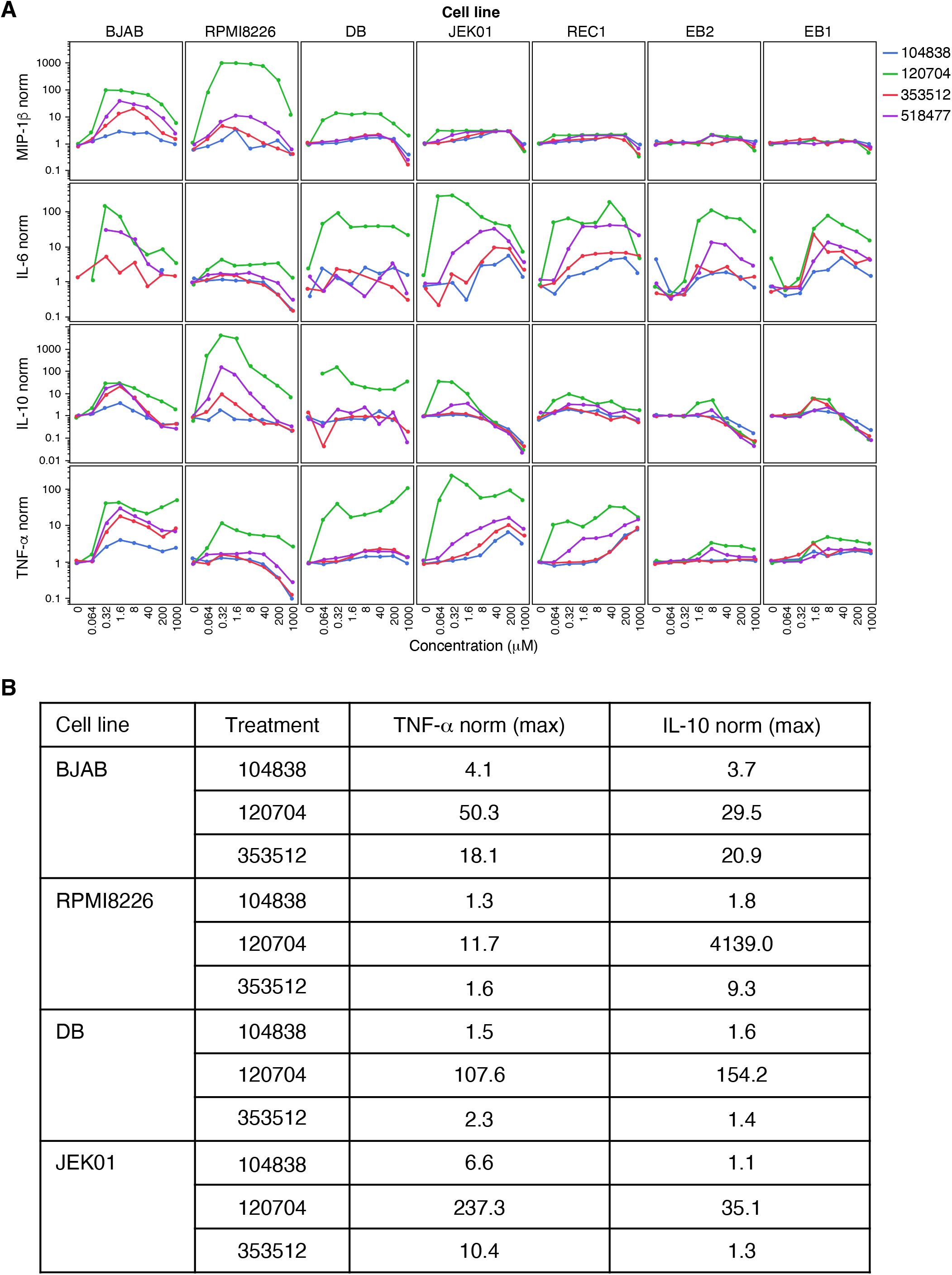
(a) MIP-1β, IL-6, IL-10 and TNF-α levels normalized to their respective untreated levels measured in the top 7 cell lines treated with 4 different oligonucleotides. (b) Maximum fold change in TNF-α and IL-10 of each oligonucleotide relative to untreated control for the top 4 discriminatory cell lines.

Importantly, Bjab cells were best capable of discriminating ISIS 353512 and the other inflammatory PS-ASOs from ISIS 104838. For example, IL-10 and TNF-α production was clearly elevated in response to ISIS 353512 as well as ISIS 518477 in contrast to treatment with ISIS 104838 which only induced minimal elevation in those analytes. The Bjab response to ISIS 353512 peaked at around 1.6 μM with an increase in TNF-α and IL-10 of 18.1 and 20.9-fold relative to untreated control, respectively, before progressively decreasing as also observed in response to treatment with ISIS 120704 or ISIS 518477 (Figure 2 B). On the other hand, ISIS 104838 induced an increase in TNF-α and IL-10 of only 4.1 and 3.7-fold relative to untreated control. In comparison, RPMI8226, DB, and JEKO1 cells show discrimination for one but not for both analytes between the benchmark PS-ASOs, and the discrimination was less dramatic. Overall, the data demonstrate the heterogeneity and complexity of TLR9 signaling pathways in response to various immune responsive PS-ASOs. The Bjab cell line, however, emerged as an encouraging potential candidate to mirror the hPBMC screening assay.

### Human PBMC and Bjab response to ASOs is highly correlated

Next, we sought to directly compare the levels of immune activation between the established hPBMC assay and the Bjab assay using a panel of 184 PS-ASOs. The ASOs had been previously tested in PBMC as part of routine screening campaigns. We compared IL-6 protein production from hPBMC cells with TNF-α protein production in Bjab cells (Figure 3A) at 3 different PS-ASO concentrations. TNF-α was chosen as it was the most robust responder in generating a wide dynamic range to control PS-ASOs (Figure 2). Analysis of the correlation between the two assays at various concentrations was performed (Figure 3B). The strongest correlation of a range of immune responsive PS-ASOs between the two assays was obtained using 0.32 μM PS-ASO administration in the PBMC assay and 1.6 μM PS-ASO administration in the Bjab assay. Interestingly this correlated with the peak of immune response in the Bjab assay. The strength of the correlation suggests that Bjab cells and PBMC cells show similar immune responses to a PS-ASOs over a wide range of PS-ASO mediated innate immunogenicity. This further establishes the more convenient Bjab assay as a suitable stand-in for the PBMC assay to evaluate the potential of PS-ASO immune responses in humans.

**Figure 3.**
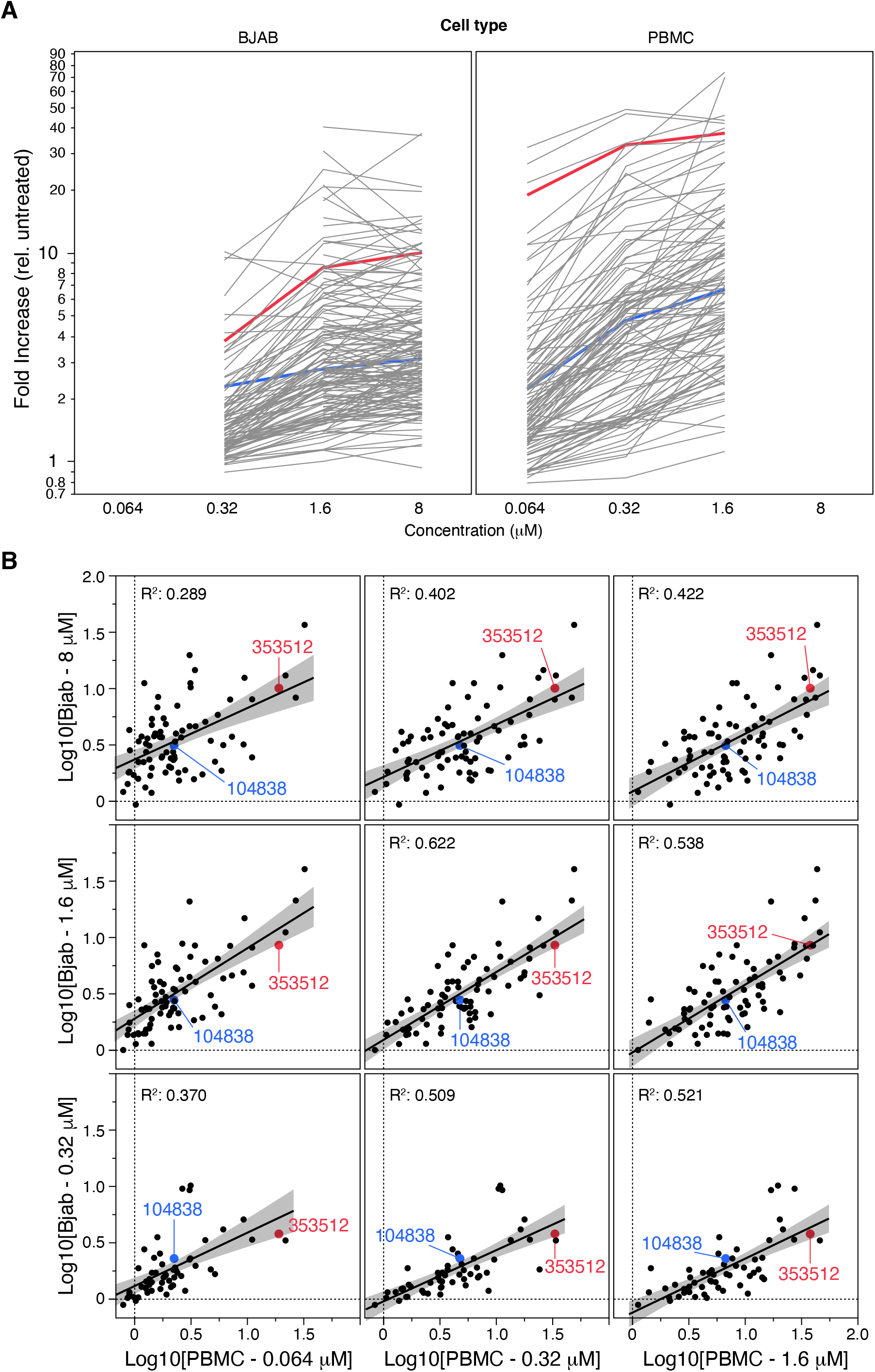
(a) TNF-α production normalized to untreated controls by Bjab cells (left), IL-6 production normalized to untreated controls by hPBMC (right) for184 ASOs. ISIS 353512 and ISIS 104838 are represented in red and blue respectively for reference. (b) PBMC IL-6 levels (X-axis) compared the Bjab TNF-a levels on the Y-axis. 3 concentrations encompassing the peak response for each cell model were compared. The correlation between hPBMC treated with ASOs at 0.32μM and Bjab treated with ASOs at 1.6 μM was highest.

### TNF-α and IL-10 stimulation by ISIS 353512 is mediated by hTLR9 in Bjab cells

The identification of a cell line responsive to ISIS 353512 in a manner reminiscent to hPBMC opened the door for the clarification of the mechanism of PS-ASO mediated cytokine response. Previous work did not conclusively establish that the activation of hPBMC cells by ISIS 353512 was mediated by TLR9 (Burel, Machemer et al. 2021). In qualifying the new assay, it was important to establish whether TLR9 was also involved in the activation of Bjab cells by non-CpG ODNs. To that end, we engineered two functionally equivalent Bjab lines in which the expression of the TLR9 was abrogated using a CRISPR based approach. Indeed, we determined that the PS-ASOs that generate an innate immune response in WT-Bjab cells, fail to do so in TLR9-KO Bjab cells (Figure 4A). This result is consistent with previous data from animal models showing the requirement of TLR9 in producing immune responses for several non-CpG ODNs (Paz, Hsiao et al. 2017). To test if the TLR9-KO cells retain functionality for other TLRs, we treated WT and TLR9-KO Bjab cells with various TLR ligands (Figure 4B). Wild-type Bjab cells Wild-type Bjab cells retained their response to the standard ligands for TLR 4, 5, and 9; the other ligands showed no responses, likely due to the lack of the functional pathways for their receptors (TLR1, TLR2, TLR3, TLR6, TLR7 and TLR8) in Bjab cells. Critically, the signal was abrogated only in the TLR9 KO cells for the TLR9 ligand, showing that the cell line maintains the capability of signaling for the other TLRs.

**Figure 4.**
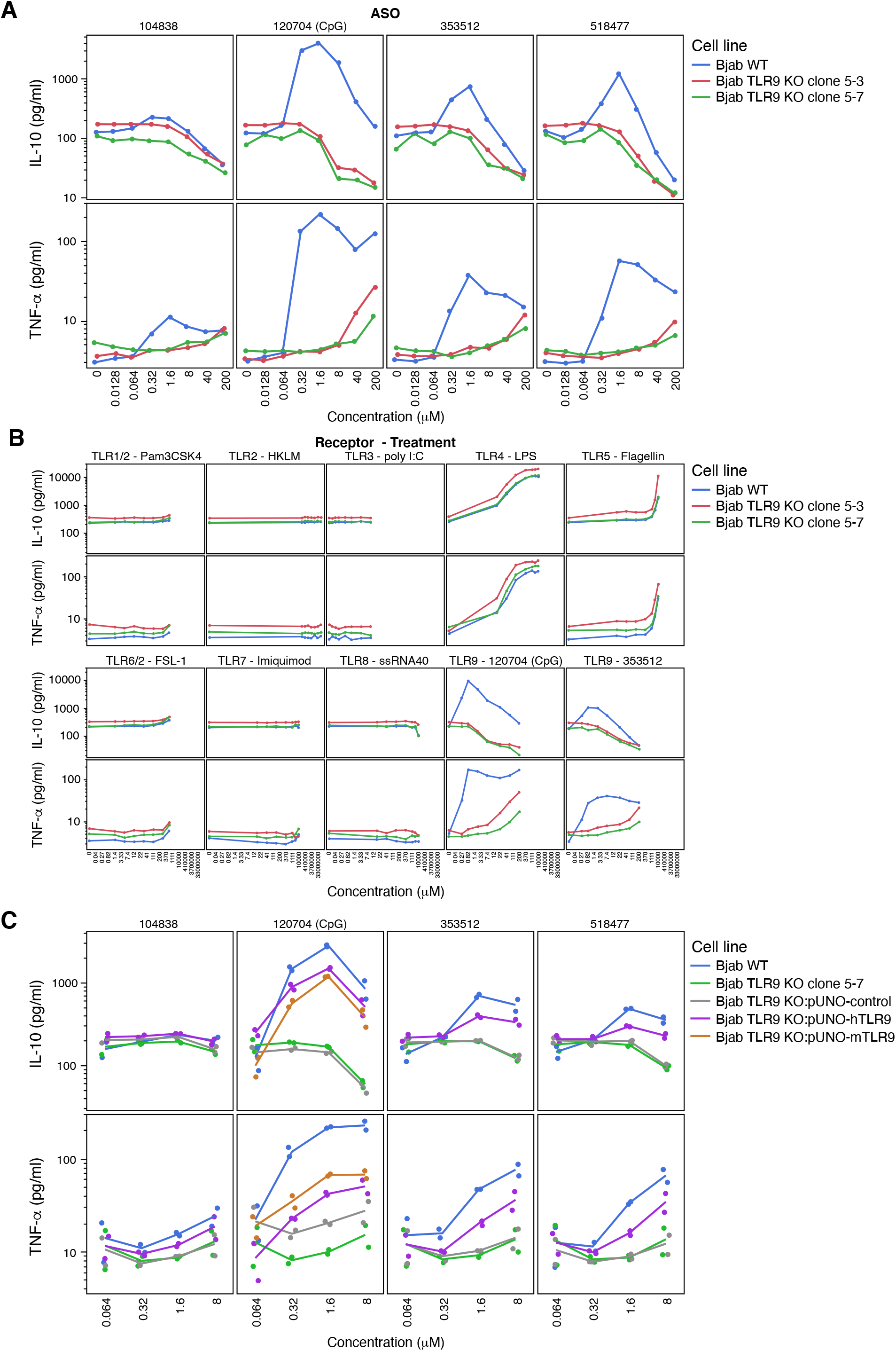
(a) Comparing Bjab response to 4 ODNs in presence of TRL9 (Bjab WT) or absence of TLR9 (Bjab TLR9 KO clone 5-3 and 5-7). (b) Comparing Bjab response to 8 different TLR antagonist in presence of TRL9 (Bjab WT) or absence of TLR9 (Bjab TLR9 KO clone 5-3 and 5-7). (c) Comparing Bjab response to 4 ODNs in presence of TRL9 (Bjab WT), absence of TLR9 (Bjab TLR9 KO clone 5-7) or with Bjab TLR9 KO reconstituted with either mouse (mTLR9) or human TLR9 (hTLR9).

Next, to further confirm the role of hTLR9 in innate immune signaling for our ODNs, we generated another series of cell line where human or mouse TLR9 was reconstituted in TLR9-KO cells (Figure 4C). The CpG ODN showed innate immune responses with both human and mouse TLR9, with the latter showing less activation, consistent with previous reports (Pohar, Yamamoto et al. 2017). However, the two non-CpG ODNs only showed responses with the human TLR9. This finding indicated that ISIS 353512 can stimulate hTLR9 but not mTLR9 at least in the context of Bjab cells. This data is consistent with previous experiments where ISIS 353512 showed negligible immune responses in mouse models yet displayed immune responses in human trials (Burel, Machemer et al. 2021). Overall, our data supports the role of human TLR9 in Bjab cells as the key mechanistic driver of human innate immune responses to various PS-ASOs and underscores the necessity of a convenient human-TLR9 based in vitro assay, which can be used for both screening purposes and mechanistic studies.

### Identification of mRNA-based markers to replace MSD-based indicator of immune activation

To further streamline this preclinical assay to determine PS-ASO based innate immune activation potential in humans, we sought to develop a much quicker RT-qPCR based mRNA analysis to replace the slower and more cumbersome MSD-based approach. We first monitored protein expression of TNF-α and IL-10 via MSD at both 6 hours and 24 hours following administration of 5 model PS-ASOs (Figure 5A). Next, we analyzed the mRNA transcripts of the same samples using 3’ end RNAseq and determined that 47 mRNA transcripts showed a high level of correlation with the extent of cytokine expression ranging from an RSquare (r^2^) of 0.323 for Shb to an r^2^ of 0.935 for Fcrl3 at 24 hours (Figure 5B, Supplemental table 1). Pathway analysis of the samples show indication of TLR pathway activation via NF-KB pathway activation, as expected (Figure 5C).

**Figure 5.**
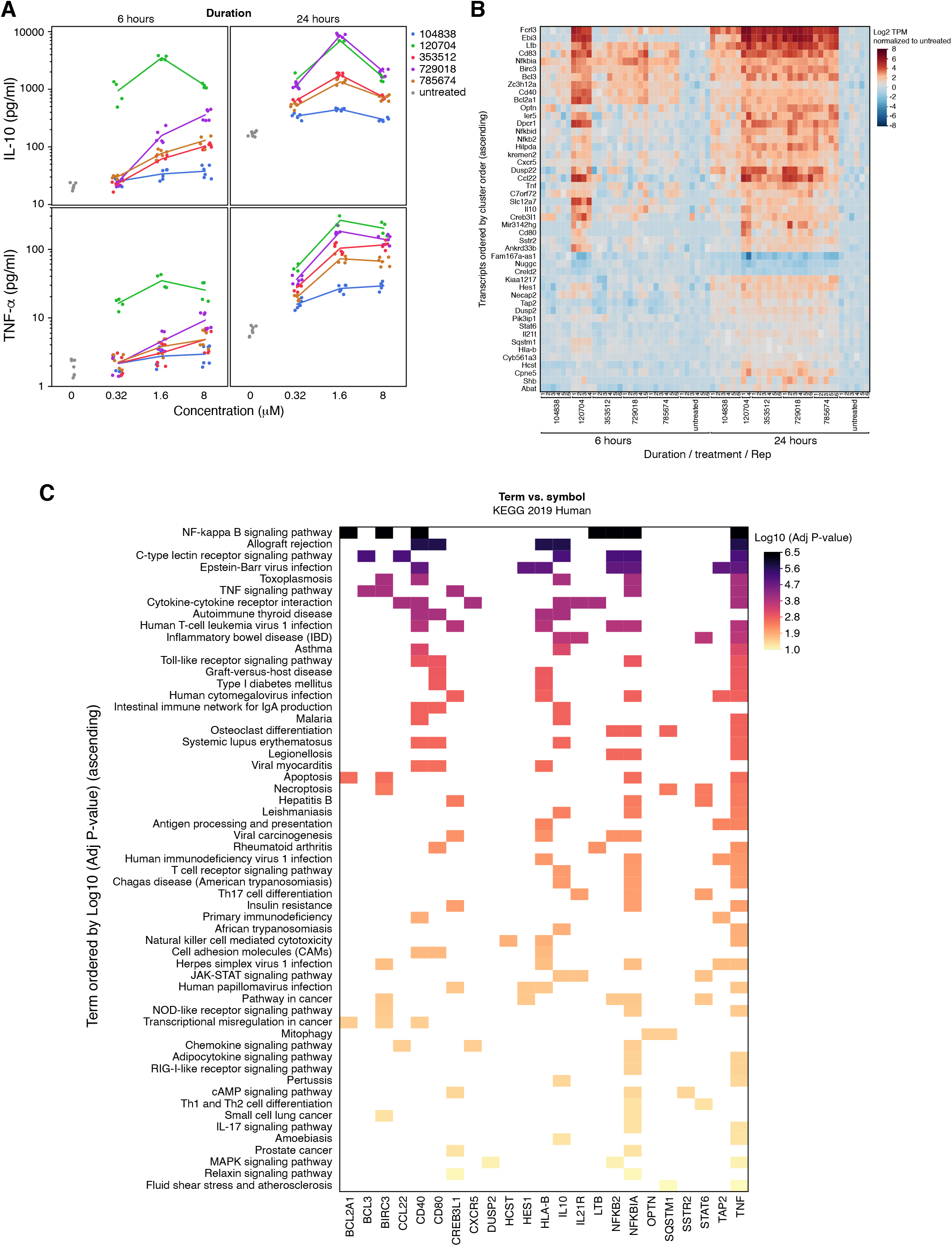
(a) IL-10 and TNF-α level measured by MSD in Bjab treated with 5 different ODN at 3 concentrations for 6 and 24 hours. (b) DGE profiling for Bjab treated with 1.6 μM ASO for 6 and 24 hours. Hierarchical clustering of the 47 transcripts most significantly correlated to TNF-α measured by MSD. Each column represents a technical replicate (c) KEGG pathway enrichment analysis of the 47 Transcripts most correlated to TNF-α. Only transcripts associated significantly enriched pathways (Y-axis) are represented.

A more detailed quantitative analysis of 4 transcripts that appeared to best mirror the MSD analysis (Ccl22, Ebi3, Fcrl3, TNF-α) indeed show high level of correlation between TNF-α protein production and mRNA transcription levels for the 5 PS-ASOs (Figure 6A), using 1.6 μM ASO at 24 hours of treatment. Next, using a panel of 66 PS-ASOs that produce varying levels of innate immune activation, we compared the MSD analysis with RT-qPCR analysis for the same 4 mRNA transcripts using the same experimental conditions (Figure 6B). Interestingly, TNF-α mRNA analysis correlated the worst with TNF-α protein production (r^2^=0.476), while Ebi3 and Ccl22 showed excellent correlation (r^2^=0.899 and r^2^=0.923 respectively) over a wide range of immune activation levels. This series of experiments demonstrates that simple and convenient RT-qPCR analysis of Ebi3 or Ccl22 mRNA performs as well as an MSD-based approach in determining the potential for innate immune activation of PS-ASOs in Bjab cells.

**Figure 6.**
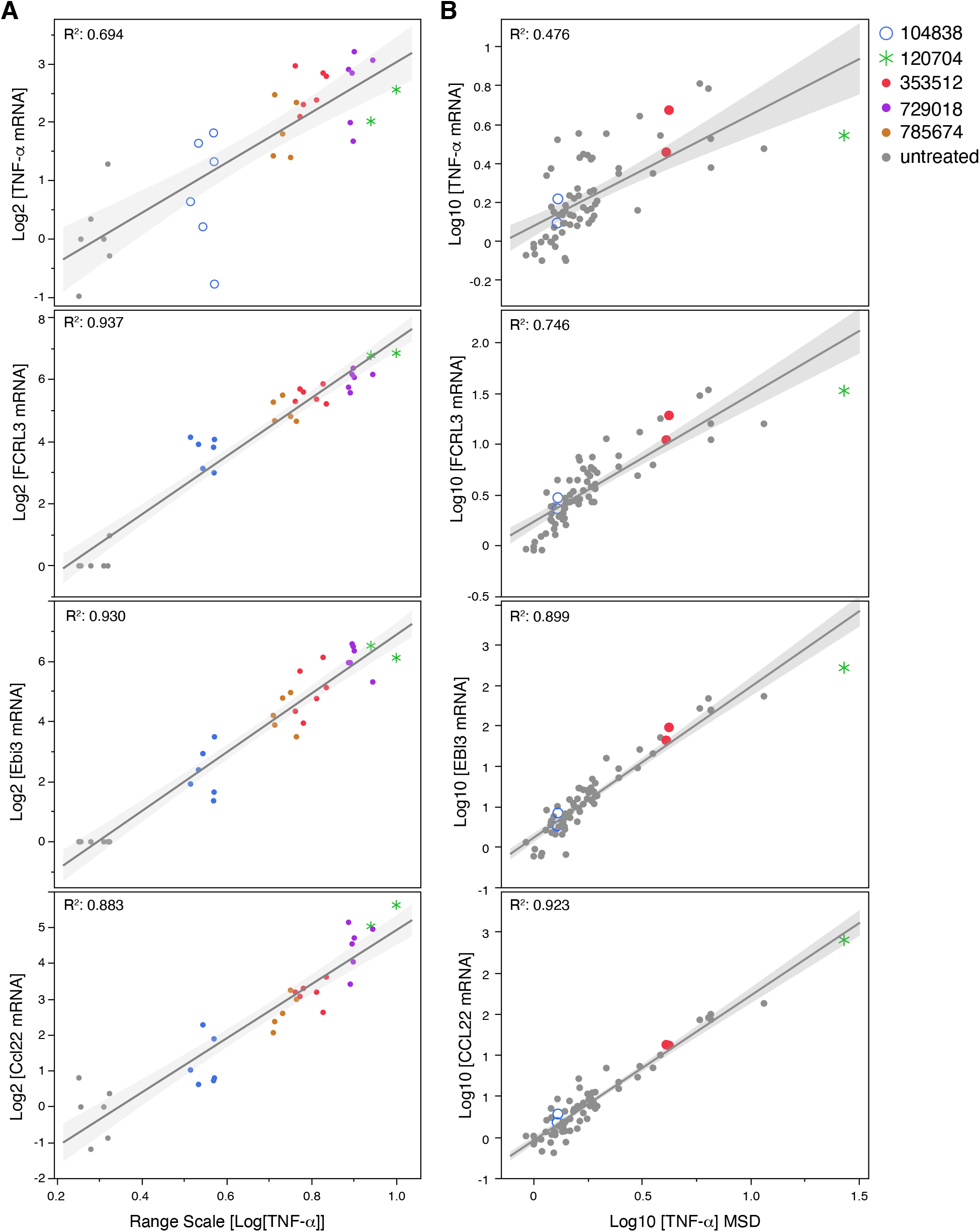
(a) Top 3 transcripts (Ccl22, Ebi3 and Fcrl3 mRNA) levels assessed by DGE (a) or RTPCR (b) from Bjab cells treated with 1.6 μM ASO for 24 hours are correlated with TNF-α protein level measured by MSD. (a) 6 replicates per ASOs were used. (b) The correlation between the novel transcripts and TNF-α protein levels was further validated with an expended set of 66 ASOs

### Bjab response to PS-ASOs is reproducible and provides a large dynamic range

The main driver for the development of an alternative assay to the hPBMC screen was to establish an assay which could be used in a higher throughput setting than the hPBMC, which requires PBMC isolated from blood collected from healthy volunteers shortly prior the assay. In contrast, an optimized cell line-based assay enables comparative ease in setting up the assay and should also be more reproducible as the cells will be genetically identical each time the assay is performed. In contrast, PBMC cells will vary due to donor variability. To that end, we monitored the performance of the assay by repeating it several times (Figure 7). Every time a screen is performed to assess the proinflammatory potential of novel PS-ASOs, Bjab cells were also treated with either ISIS 104838 or ISIS 353512 as benchmark controls of low and high innate immune activation respectively. In addition, ISIS 735746 and ISIS 785674 are included as intermediate markers of inflammation. By performing these controls on multiple occasions, the investigator can assess the quality of the assay and its ability to identify and exclude new inflammatory PS-ASOs.

**Figure 7.**
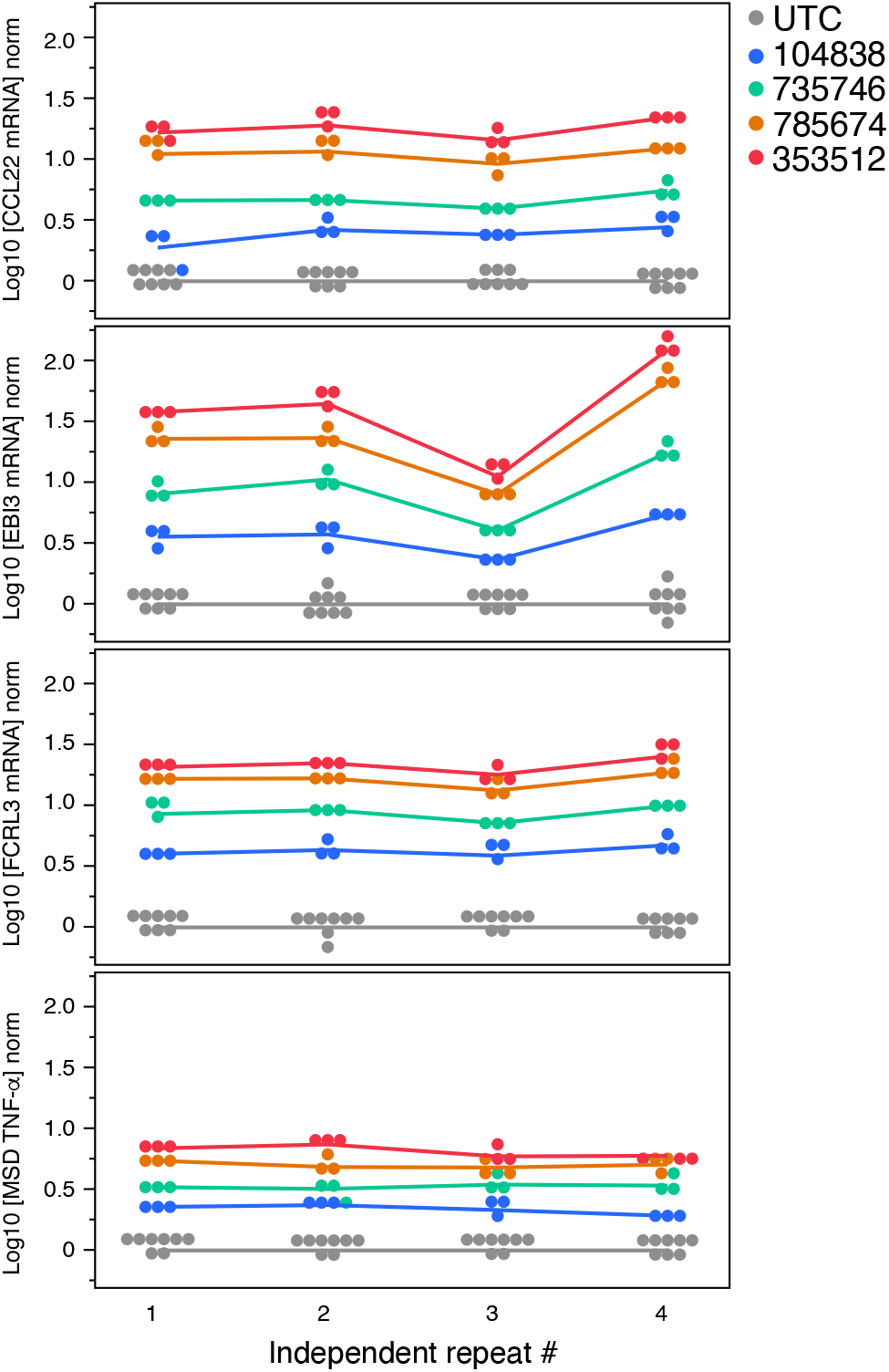
Assessing reproducibility. 4 independent runs were conducted comparing TNF-α protein levels as measured by MSD with Ccl22, Fcrl3 and EBI3 mRNA levels measured by RT-PCR

Across multiple experiments, a clear discrepancy in immune responses between the PS-ASOs was observed with remarkable consistency. This affords the investigator ease in using statistical analysis in determining the precise inflammatory potential of certain PS-ASOs. While the three mRNA markers all performed well, Ccl22 showed the most consistent data across multiple separate experiments and had the largest dynamic range between low and high levels of PS-ASO innate immune activation. While TNF-α protein levels demonstrated a clear dynamic range between the benchmark ASOs, the discrepancy is not as stark as the mRNA analysis. Thus, Ccl22 mRNA analysis was chosen as the most robust responder for the assay.

Prior to this analysis, assessment of the Bjab assay benchmark controls (ISIS 104838 or ISIS 353512) using TNF-α protein production as a readout was performed during PS-ASO screening over a several year period (Supplemental Figure S2). On some occasions, the dynamic range between ISIS 104838 and ISIS 353512 was too narrow to confidently identify PS-ASOs with increased proinflammatory potential, and the screen was repeated. However, even in this context, a suitably large dynamic range was apparent, further demonstrating the robustness of the Bjab assay, even when an unoptimized readout was employed.

## DISCUSSION

PS-ASOs have been proven to be safe and potent drugs across various therapeutic areas. In select cases, however, some PS-ASOs can cause unwanted effects, including cytotoxicity (Shen, De Hoyos et al. 2019) and innate immune activation (Frazier 2015). Ideally, these PS-ASOs with stronger proinflammatory potential can be identified and discarded early in the developmental process. Ongoing efforts to understand PS-ASO behavior and subsequent improvement of PS-ASO technology will aid in this and improve PS-ASO developmental processes. ISIS 353512 caused pro-inflammatory indications in healthy human volunteers while pre-clinical animal models showed no indication of such effects. This highlights the species-specific nature of these processes and underscores the need for an in vitro human-based cell line to for pre-clinical screening. While the hPBMC assay provides this, it suffers from donor-to-donor variability and is a resource-intensive assay.

To identify a suitable cell line that would be responsive to ODNs in a manner that replicates the HPBMC assay, we first queried two databases cataloging TLR9 expression levels. Based on previous data linking PS-ASO immune activation with the TLR9 pathway, we anticipated that TLR9 levels would be correlated with immune activation and that trends for the CpG and non-CpG ODNs would be similar. This was generally not the case. Most cell lines tested herein showed responses to the CpG ODNs, and the CpG ODNs was considerably more potent in comparison to the non-CpG ODNs, consistent with previous data (Younis, Vickers et al. 2006). However, some of the cell lines (Karpas422 and MEC2) showed no response to the CpG ODN, in spite of the presumed presence of a functional TLR9 pathway. The CpG ODN also showed slightly more consistent readout of the immune markers within each cell line in comparison to the non-CpG ODNs. For example, IL-6 and TNF-α show drastically different expression profiles within each cell line in response to the non CpG ODNs. Overall, analysis of several cytokines and chemokines in 14 of the most promising cell lines showed drastically varied results, leaving little opportunity to identify meaningful patterns or trends. This is particularly well illustrated for the non-CpG ODNs. One potential explanation for this phenomenon lies in TLR9’s ability to bind to various ODNs, regardless of their activating potential. This can result in full, partial, or no TLR9 pathway activation, and may result in the activation of only certain inflammatory markers (Ohto, Ishida et al. 2018).

Some Non-CpG ODNs appear to be being able to activate TLR9, but are relatively poor substrates and may not give full pathway activation relative to CpG ODN. This is more apparent in some cell lines compared to others where the ability of those cell to active and signal through TLR9 might be compromised at various steps of the TLR9 signaling cascade. This highlights the complexity of PS-ASOs immune activation via non-CpG ODNs and shows that much work is needed to fully understand these mechanisms.

The model PS-ASOs that have been shown to display immune reposes in the PBMC model, or in clinical trials, showed no responses in many of the cell lines used here. Importantly, however, investigation of this panel of cells produced one especially promising candidate, the Bjab cell line, where a clear discrepancy was determined between benchmark PS-ASOs ISIS 104838 and ISIS 353512 for both IL-10 and TNF-α protein production. Interestingly, a previous report suggests the role of Bjab cells as responsive to TLR9 agonists (Noack, Jordi et al. 2012). Importantly, the Bjab assay is reliably able to assess the immune potential of PS-ASOs over a wide range of immune responses in a manner that corresponds with the hBPMCs. Next, the Bjab assay was further streamlined to replace the time and resource intensive MSD-based analysis with the more convenient RT-qPCR based analysis, with Ccl22 mRNA production identified as the best predictor of TNF-α protein production. Finally, analysis of benchmark ODNs over several independent trials of performing this assay showed robust and consistent differences in activation. This underscores an advantage to the Bjab assay in that the HPBC assay suffers due to variability from different donors, resulting in a lack of consistency when repeating the assay.

The Bjab assay not only acts as a screening tool, but also can also provide critical mechanistic insights. Here, we provide additional evidence that ODNs such as ISIS 353512 can activate TLR9 in a species-specific fashion thereby corroborating exhaustive pre-clinical and clinical efforts where ISIS 353512 showed no immune activation in animal models but caused immune activation in humans (Szalai, McCrory et al. 2014, Burel, Machemer et al. 2021). With the Bjab assay now in place, such information can be gained with minimal effort. Furthermore, our analysis underscores the discrepancy between human and mouse TLR9 pathway activation, positioning the Bjab cells as an excellent pre-clinical model for studies of cellular TLR9 pathway activation. For example, diseases involving non-CpG nucleic acid mediated TLR9 activation (Garcia-Martinez, Santoro et al. 2016), such as in certain auto-immune diseases (Gilliet, Cao et al. 2008), could be studied in this model as opposed to mouse models or other animal cell lines. Overall, the Bjab assay is a robust pre-clinical tool that replaces the hPBMC assay as an indicator of human PS-ASO innate immune activation.

## ACKNOWLEGMENTS

The authors thank the donors who participated in this study; Hongda Li for her help in critical review of an earlier version of the article; and Michael Cichanowski for assistance with formatting the figures and Angela Colabucci and Tamika Holmes in proofreading and preparing the manuscript for publication.

