## supplemental figures and table for "Mechanism Driven Early Stage Identification and Avoidance of Antisense Oligonucleotides Causing TRL9 Mediated Inflammatory Responses in Bjab cells"

**Supplemental Figure 1**

7 additional cell lines were treated with 4 ODNs and MIP-1β, IL-6, IL-10 and TNF-α levels were measured by MSD. Normalized values relative to their respective untreated levels are shown.

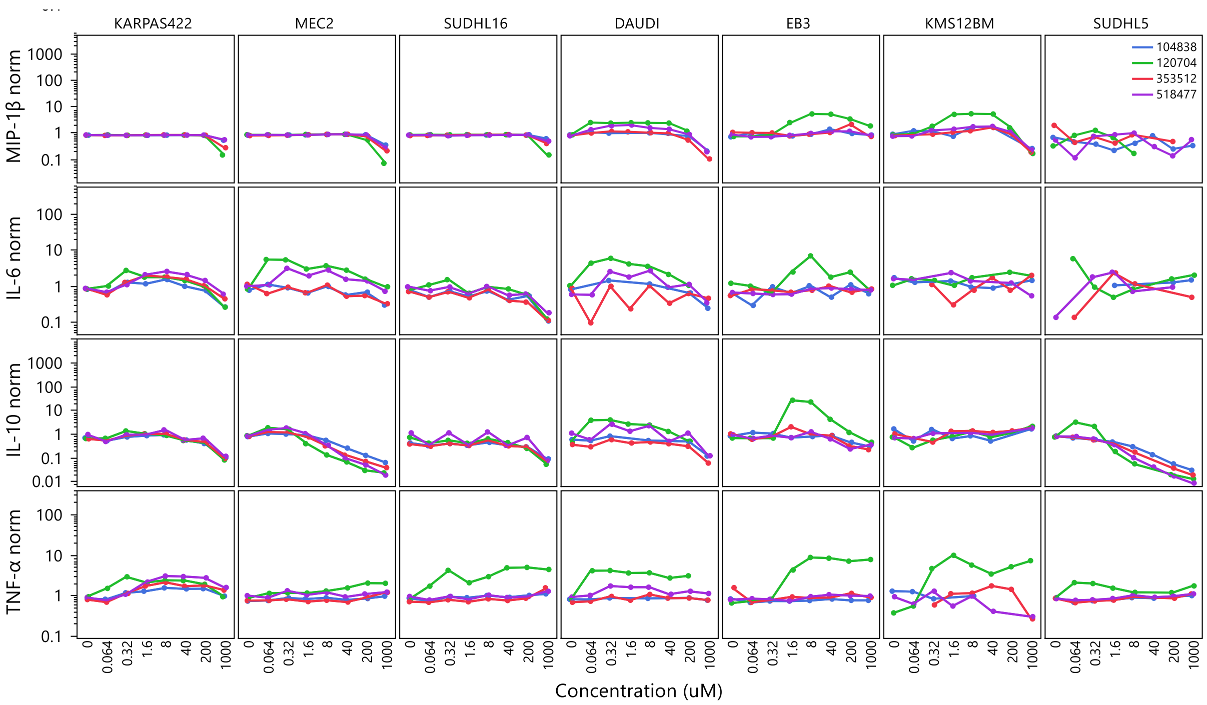

**Supplemental Figure 2**

Range of relative changes in TNF-α production in Bjab following treatment with either ISIS 104838 (open circles) or ISIS 353512 (closed circles). Results from 22 independent experiments performed on different date are represented separately.

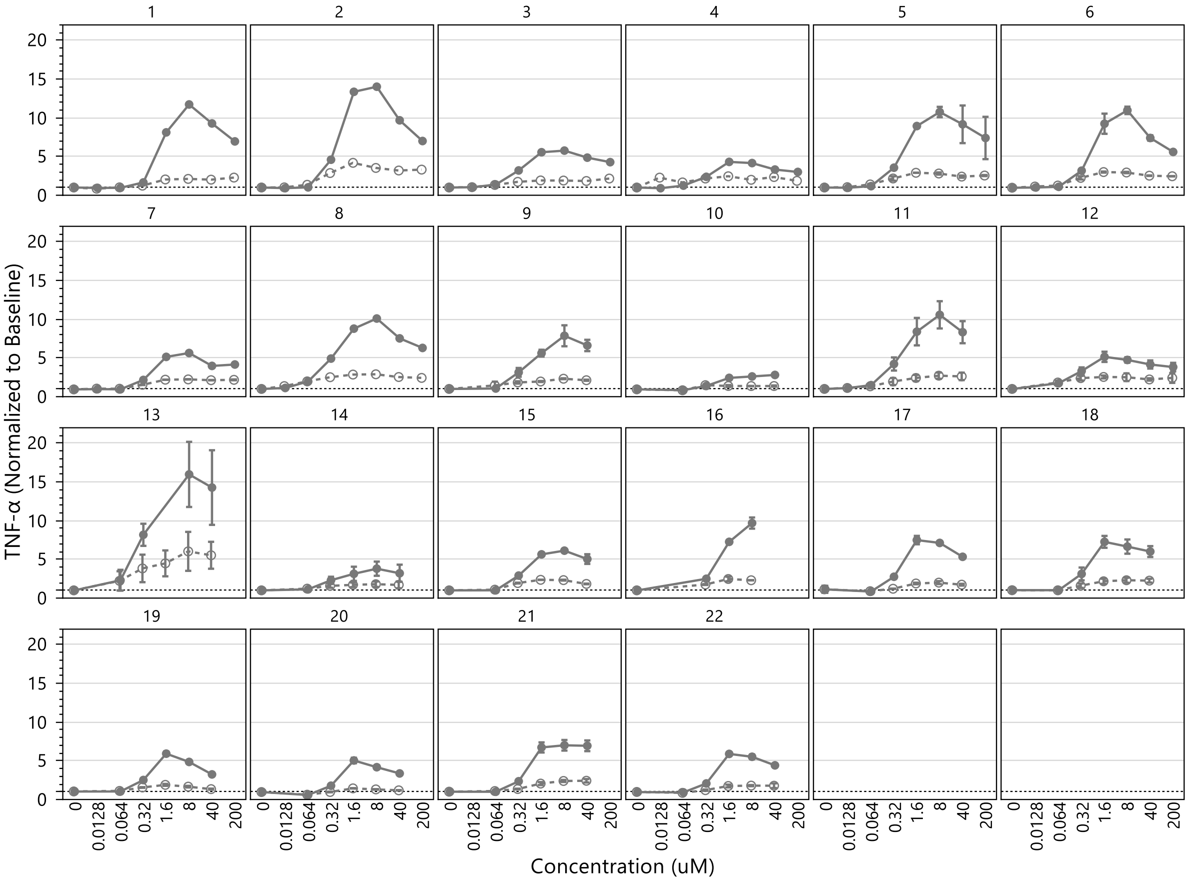

**Supplemental table 1**

Rsquare for the top transcripts measured by DGE correlated to TNF-α protein level measured by MSD

| | RSquare Adjusted | |
|---|---|---|
| **Symbol** | **06 hrs** | **24 hrs** |
| Fcrl3 | 0.687 | 0.935 |
| Ebi3 | 0.782 | 0.927 |
| Cd83 | 0.582 | 0.891 |
| Ccl22 | 0.476 | 0.879 |
| Bcl2a1 | 0.764 | 0.843 |
| Birc3 | 0.524 | 0.838 |
| Cd40 | 0.697 | 0.819 |
| Nfkbia | 0.233 | 0.816 |
| Sstr2 | 0.223 | 0.799 |
| Dpcr1 | 0.730 | 0.770 |
| Ankrd33b | 0.504 | 0.764 |
| Zc3h12a | 0.513 | 0.764 |
| Ltb | 0.572 | 0.763 |
| Hilpda | 0.558 | 0.755 |
| Nfkbid | 0.263 | 0.750 |
| Il10 | 0.542 | 0.730 |
| Cpne5 | 0.065 | 0.721 |
| Optn | -0.031 | 0.712 |
| Creb3l1 | 0.32 | 0.703 |
| Dusp22 | 0.465 | 0.695 |
| Tnf | 0.35 | 0.684 |
| C7orf72 | 0.344 | 0.670 |
| Nuggc | 0.400 | 0.654 |
| Necap2 | 0.352 | 0.652 |
| Nfkb2 | 0.375 | 0.645 |
| Bcl3 | 0.417 | 0.638 |
| Hes1 | 0.506 | 0.635 |
| Kremen2 | 0.577 | 0.632 |
| Cxcr5 | 0.379 | 0.63 |
| Sqstm1 | 0.700 | 0.629 |
| Kiaa1217 | 0.072 | 0.606 |
| Cyb561a3 | 0.341 | 0.605 |
| Mir3142hg | 0.656 | 0.593 |
| Pik3ip1 | 0.039 | 0.582 |
| Dusp2 | 0.319 | 0.575 |
| Ier5 | 0.408 | 0.567 |
| Hcst | -0.024 | 0.551 |
| Creld2 | 0.148 | 0.518 |
| Slc12a7 | 0.717 | 0.503 |
| Hla-b | 0.681 | 0.476 |
| Fam167a-as1 | 0.497 | 0.453 |
| Cd80 | 0.443 | 0.450 |
| Stat6 | 0.212 | 0.403 |
| Tap2 | 0.001 | 0.391 |
| Abat | 0.17 | 0.363 |
| Il21r | 0.192 | 0.355 |
| Shb | 0.429 | 0.323 |
